# IVT generation of guideRNAs for Cas9-enrichment Nanopore Sequencing

**DOI:** 10.1101/2023.02.07.527484

**Authors:** Timothy Gilpatrick, Josh Zhiyong Wang, David Weiss, Alexis L Norris, James Eshleman, Winston Timp

**Affiliations:** Department of Biomedical Engineering, Johns Hopkins University, Baltimore, MD, USA; Agilent Technologies, Santa Clara, California, USA; Departments of Pathology and Oncology, The Sol Goldman Pancreatic Cancer Research Center, Johns Hopkins School of Medicine, Baltimore, MD, USA; Department of Molecular Biology and Genetics, Department of Medicine, Division of Infectious Disease, Johns Hopkins School of Medicine, Baltimore, MD, USA

## Abstract

Generating high-coverage sequencing coverage at select genomic loci has extensive applications in both research science and genetic medicine. Long-read sequencing technologies (e.g. nanopore sequencing) have expanded our ability to generate sequencing data in regions (e.g. repetitive elements) that are difficult to interrogate with short-read sequencing methods. In work presented here, we expand on our previous work using CRISPR/Cas9 for targeted nanopore sequencing by using *in vitro* transcribed guideRNAs, with 1100 guideRNAs in a single experiment. This approach decreases the cost per guideRNA, increases the number of guideRNAs that can be multiplexed in a single experiment, and provides a way to rapidly screen numerous guideRNAs for cutting efficiency. We apply this strategy in multiple patient-derived pancreatic cancer cell lines, demonstrating its ability to unveil structural variation in “deletion hotspots” around the tumor suppressor genes *p16* (*CDKN2A*), and *SMAD4*.

## INTRODUCTION

Sequencing of nucleic acids has revealed incredible insight into living organisms, opening up fields of investigation and revolutionizing medicine and biology. The human genome with its 3 billion nucleotides has finally been assembled tip-to-tip^1^. However, for many biological questions it is not necessary to sequence the entire genome, instead targeted sequencing can reduce the cost of generating and analyzing sequencing data. This has motivated the development of strategies for loci enrichment, but most strategies are tailored to short-read sequencing platforms. This motivates the development of additional tools for target enrichment that are more compatible with long-read sequencing.

Our group previously described on-target Cas9 enrichment to generate high coverage at regions of interest with nanopore sequencing^2^. For initial studies, we evaluated multiple features - single nucleotide changes, DNA methylation, and structural variation (SV). We simultaneously interrogated ten sites with sizes 10-30kb, using a pool of ∼50 guideRNAs (gRNAs). For those studies, it was necessary to know *a priori* the location of genomic aberrations being queried with relative precision. This type of targeting can introduce bias during the discovery of novel breakpoints, as frequently the sites of structural variations may not be known. For instance in pancreatic ductal adenocarcinoma (PDAC) “hotspots” for SVs exist where the breakpoints of these aberrations occur over multiple megabases.

Two genomic regions of interest were selected for this study. Both contain tumor suppressor genes commonly inactivated through deletions in pancreatic cancer and associated with poor outcomes: *p16* (*CDKN2A*), and *SMAD4*^3–5^. To survey the large region known to contain SV breakpoints, we targeted 9Mb centered around *p16* and 5.4Mb centered around SMAD4 for a total targeted region of 14.4Mb (see Methods). The gRNAs were tiled across this 14.4Mb region, with the goal of having one gRNA every 10kb.

To avoid the high cost of synthetic gRNAs, we generated gRNAs via *in vitro* transcription (IVT), from a pool of PCR-amplified gRNA templates and adapted the nanopore Cas9-enrichment strategy for use with this large library of gRNAs. We first demonstrated the method’s efficacy using the model cell line GM12878, and then applied the strategy to three patient-derived PDAC cell lines. SVs were determined from the targeted nanopore data, and results compared with SVs previously identified using SNP (Illumina Omni2.5M) data which had generated imprecise breakpoints and initial characterization of the SVs^3 6^.

In addition to reducing cost and increasing the number of regions that can be targeted in a single sequencing run, this IVT gRNA strategy offers a rapid high throughput method for evaluating the cutting efficiency of gRNAs. Numerous in silico tools exist for predicting the cutting efficiency of gRNAs using intrinsic sequence-specific features ^(Doench et al. 2016; Farboud and Meyer 2015; Chari et al. 2015)^, and the ability to predict gRNA cutting efficiency continues to be an active area of investigation^10^. Despite these efforts, many gRNAs with good “on-target” scores show great variability in their efficiency at inducing DNA cleavage. Therefore, a strategy for rapid gRNA evaluation has the potential to streamline CRISPR/Cas9 workflows with new guideRNAs and can be used to help improve gRNA prediction tools.

## RESULTS

### On-target Enrichment

The Cas9-enrichment strategy previously described^2^ was adapted for use with IVT gRNAs (Figure 1A, see methods). 1,104 gRNA probes were selected for the final design based on probe scores (Doench Score^7^ and Zhang Score^11^) and probe location. All gRNAs were on the positive sense strand of the DNA, with an average spacing between gRNAs of 13kb (85% between 5-21kb, largest of 42kb) (Figure 1B).

**Figure 1.**
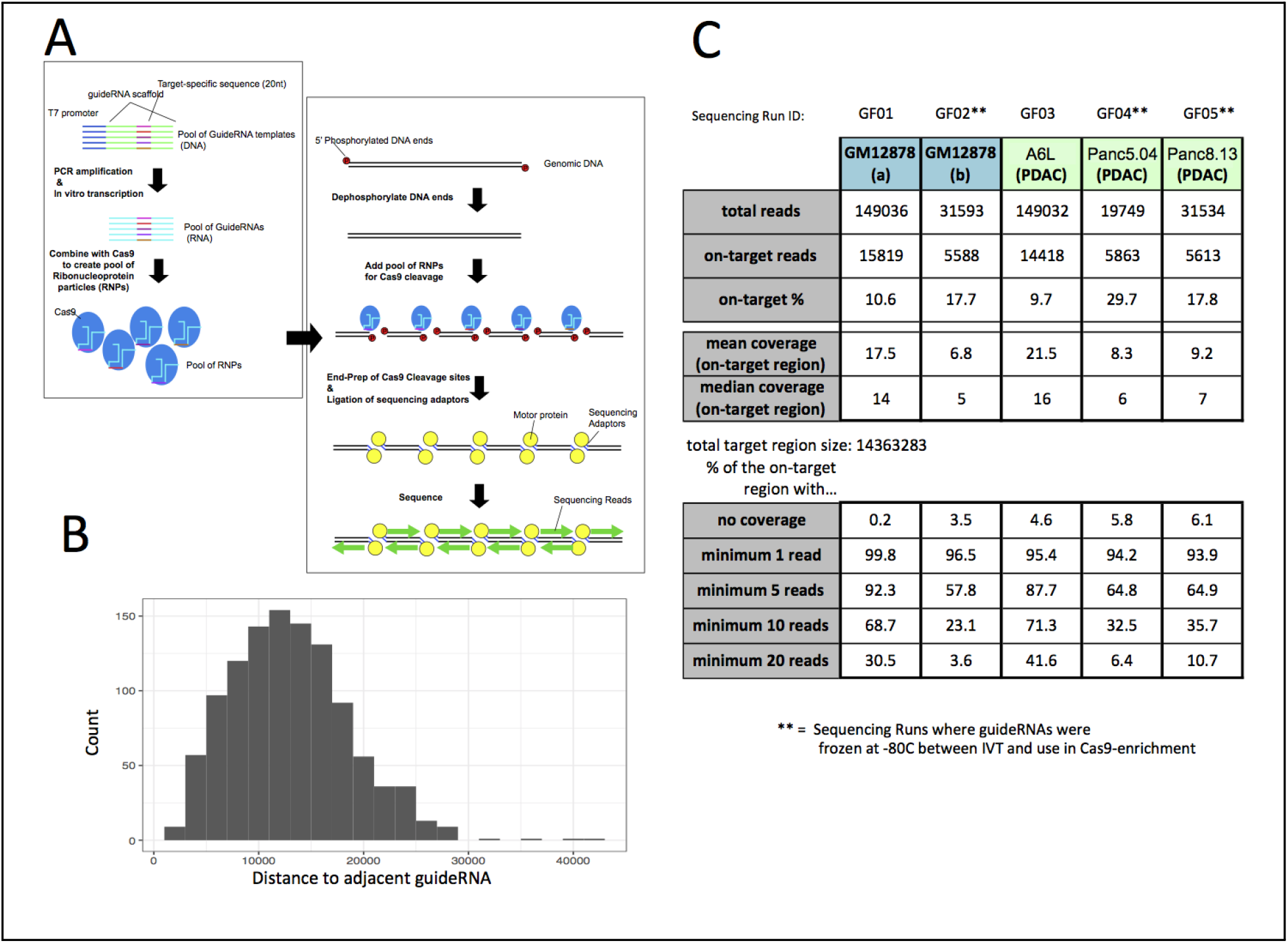
Overview of approach, guideRNA spacing, and sequencing run metrics. A. Cartoon showing the strategy for generating IVT gRNAs. B. Histogram showing the distance between gRNAs. C. Read count and on-target enrichment values for each of the sequencing runs.

The template DNA containing gRNA template sequence and T7 promoter was amplified by PCR, then used for IVT generation of gRNAs. For sequencing run IDs GF01 and GF03, the IVT gRNAs were used immediately after preparation. These runs generated ∼150,000 sequencing reads each with approximately 10% of the generated reads being “on-target” (Figure 1C). For three sequencing runs (GF02, GF04, and GF05) the gRNAs were stored at -80°C after transcription and before use in enrichment. These sequencing runs generated 20,000-30,000 reads per run (about 5-fold reduction), albeit with a slightly higher percentage of “on-target” reads (Figure 1C). The targeted region showed a wide distribution in coverage (Figure 1C, Supplementary Figures 1-2) but importantly, only very rare locations (0.2 - 6.1%) had no reads present (Figure 1C).

### Identification of Structural Variants

Three PDAC cell lines were evaluated using this approach (A6L, Panc5.04, and Panc8.13). SNP array data had previously identified the SV breakpoints for each cell line, albeit with imprecise calls^3^ (Figure 2A). Many PDAC cell lines have been further evaluated and identified the “deletion” as the result of composite rearrangements (specifically, a translocation on one side of the gene paired with an inversion on the other side)^6^. To identify SVs using our targeted nanopore data we used the Sniffles variant caller (v2)^12,13^. The variants identified included previously identified (Figure 2, Supplementary Figure 3) but also previously unannotated SVs in these cell lines (Supplementary Table T1).

**Figure 2.**
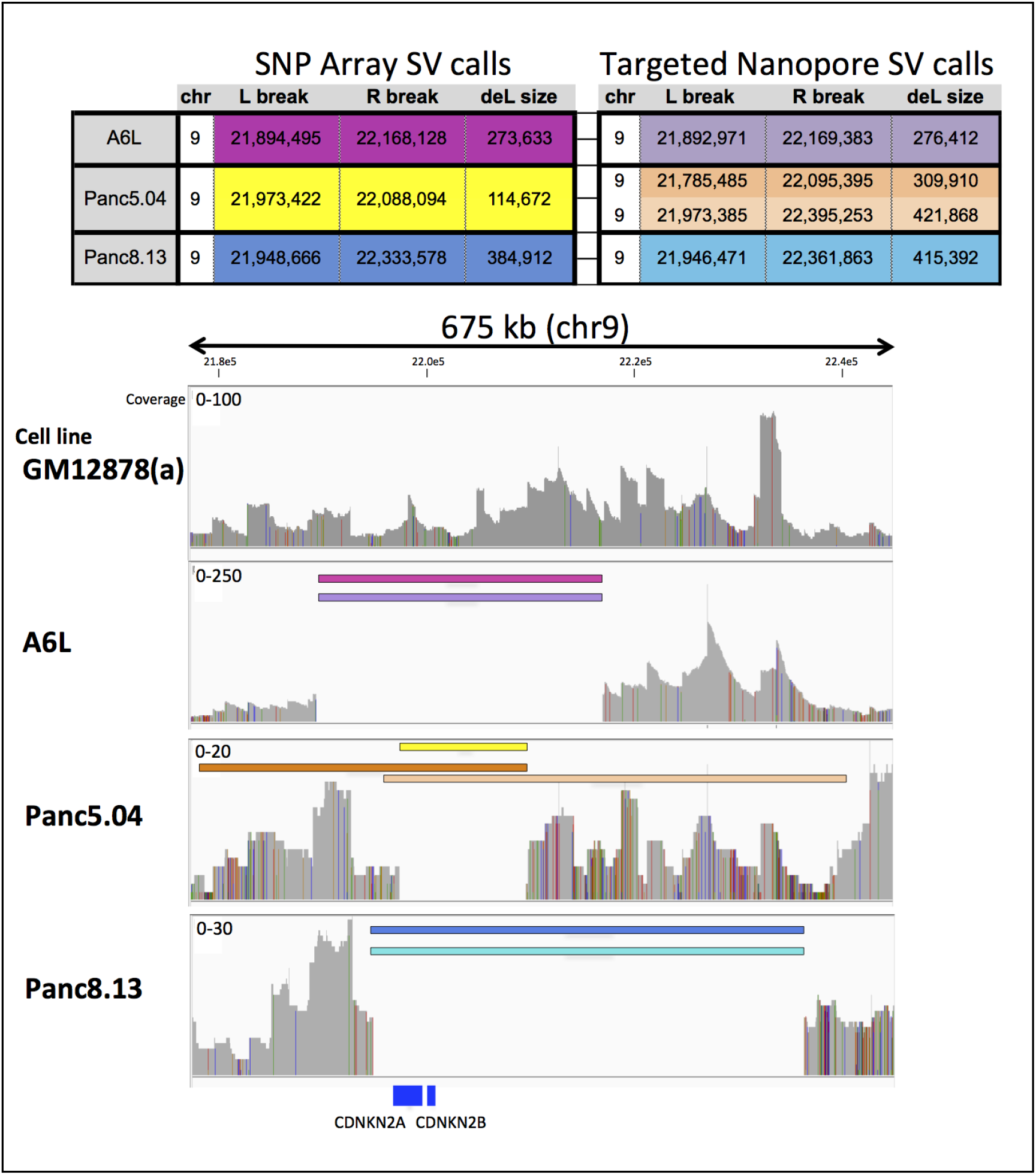
Comparing SV calls and coverage plots in GM12878 and PDAC cell lines. Top: Table comparing SV calls in the three PDAC cell lines (Omni 2.5 SNP array data^3^ compared with Sniffles v2 calls on Cas9-enrichment nanopore data). Bottom: Coverage plots across the enriched regions, showing the large deletions called. A6L and Panc8.13 have homozygous deletions. The deletion previously called as homozygous in Panc5.04 (yellow bar) is produced from two overlapping large deletions (orange bars).

For the A6L and Panc8.13 cell lines, the large previously described deletions (on chr9 and chr18) were identified by Sniffles, with homozygous breakpoints identified within 10kb of those found by SNP array. For the Panc5.04 cell line, the 460kb deletion previously annotated on chr18 (flanking *SMAD4*) was identified by Sniffles, however, the 115kb deletion on chr9 was unveiled to be a composite of two much larger overlapping heterozygous deletions (Figure 2). In addition to these noted deletions, inspection of all SV calls generated by Sniffles showed rich complexity in the targeted regions, identifying additional large deletions as well as multiple translocations (Supplementary Table T1).

### Evaluation of sgRNA Probe Performance

In addition to tiling across large regions - the ability to multiplex greater than one thousand gRNAs allows us to empirically test gRNA performance. During the design process, a weighted composite of the Doench score^7^ and Zhang score^11^ was used for initial selection (Figure 3A). As previously noted, we observed a great variability in the number of reads generated by each gRNA. Because a gRNA with low cutting efficiency could have coverage in its “on-target” region generated by cleavage of an adjacent gRNA, we used the number of reads *starting* adjacent to each gRNA (rather than coverage) to evaluate cutting efficiency. For this analysis, we averaged the two sequencing runs from GM12878. Figure 3A shows a histogram of reads generated by each gRNA, demonstrating that the large majority of the gRNAs performed poorly (<2 reads per gRNA), with a much smaller number of high-performing gRNAs (maximum of 40 reads per gRNA).

**Figure 3.**
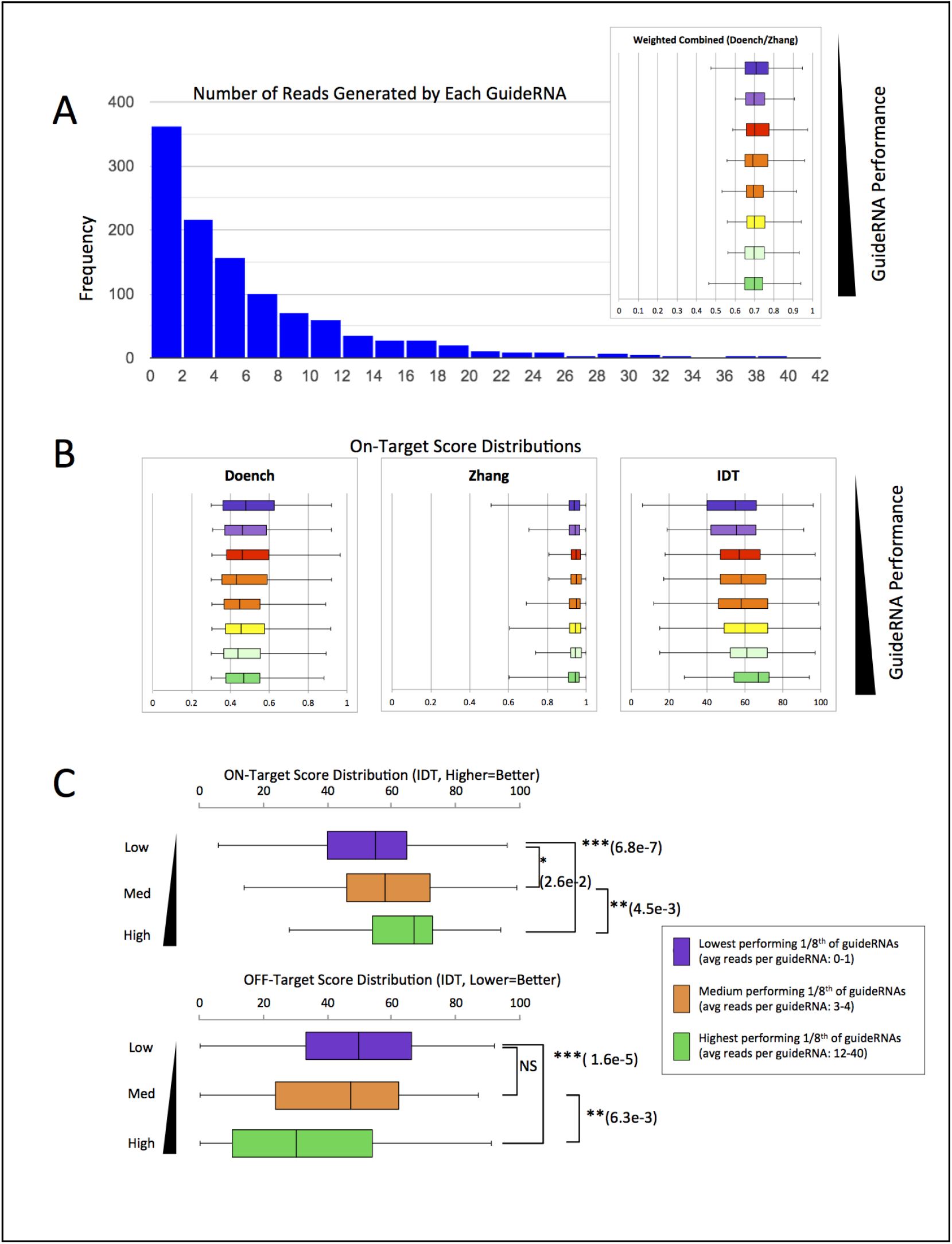
GuideRNA performance versus predicted scores using GM12878 cell line data. A. Histogram showing the average read count generated by all gRNAs (average of two GM12878 runs), and distribution of the weighted Doench and Zhang scores for the gRNAs, split into eight groups based on performance. B. Comparison of on-target scores called by multiple metrics in gRNA groups. C. Comparison of IDT on-target and off-target scores lowest, middle, and highest 1/8th of gRNAs. Statistical significance and p-value shown is using a two-tailed t-test to compare groups.

To evaluate differences in the predicted on-target performance and predicted off-target sites of gRNAs, the gRNAs were split into eight groups, ranked by the number of reads generated (Figure 3). We compared the predicted on-target score for Doench, Zhang, and the IDT online tool^14^ versus gRNA performance in these different groups. Of these three scores, only the IDT score showed correlation between cutting efficiency and score (Figure 3B). The lowest performing 1/8th of gRNAs, the highest performing 1/8th of gRNAs, and the central performing 1/8th of gRNAs were plotted to compare both on-target and off-target scores from IDT (Figure 3C). There was a large amount of variability evident in all groups, with fewer low on-target scoring gRNAs in the high-performing group. Conversely, for off-target scores, although there was a correlation between lower (better) scores and cutting, even the high-performance group had many high (worse) off-target scores, indicating off-target Cas9 cleavage had less impact.

## DISCUSSION

The work represented herein offers an initial foray into using massively multiplexed *in vitro* transcribed guideRNAs for Cas9-enrichment with nanopore sequencing. We recognize that there is substantial room for improvement and optimization, but feel that this strategy has enough important implications to merit sharing with the scientific community. This work shows that ≥1,000 gRNAs can be cheaply generated and used in a single experiment with the Cas9-enrichment nanopore protocol. This can be adapted to interrogate very large regions, as we have done here, to: (1) target large numbers of sites anticipated to be clinically informative or (2) evaluate many gRNAs simultaneously.

We recognize several shortcomings with the gRNA set being used for this initial work. Firstly, the vastly differing performance of the gRNAs suggests that optimization in gRNA selection could be done to identify higher-performing gRNAs and help achieve a more balanced coverage across the entire area being targeted. Secondly, there is likely further improvement in the spacing of the gRNAs that could be achieved. We initially sought to have the guideRNAs spaced ∼10kb, but given the long-read nature of nanopore sequencing, many reads span the entire distance between gRNAs. Despite these limitations, our application demonstrates the power of this approach to evaluate complex SVs present in cancer samples.

We also offered an analysis of how the varying gRNAs performed relative to predicted on-target cutting efficiency. The gRNA performance showed correlation with predicted on-target guideRNA cutting efficiency with IDT scores only while noting that all guides used had initially passed a minimum threshold with the Doench and Zhang scores. A noteworthy limitation of our analysis is that gRNAs were designed against the reference sequence (hg19), not considering mutations within the cell lines. Both the data generated here by our group and future implementations of this strategy, offer a dynamic way to interrogate Cas9 cutting efficiency, helping to build *in vitro* datasets to help guide our understanding of the features that lead to efficient cleavage by the Cas9/gRNA complex.

## MATERIALS & METHODS

### Guide RNA design

The gRNA sequences were designed at large loci flanking two genes of interest (*SMAD4* - chr18:46,137,962-51,507,587 and *p16* - chr9:19,631,719-28,644,497 [hg19]).The gRNAs were designed using Agilent SureDesign gRNA design software, based on two scoring algorithms with Doench Score setting at 0.3 and Zhang Score setting at 0.5 with equal weight^7,11^. This identified 3,600 candidate gRNAs sequences – 1,104 were selected to maintain ∼8-25kb spacing between adjacent gRNAs, with all gRNAs on the forward strand.

For each gRNA, the 20nt complementary sequence was inserted into the gRNA template, and a T7 promoter (5’-TAATACGACTCACTATA-3’) was added to the beginning of the sequence. The gRNA DNA templates were produced using Agilent SurePrint oligo synthesis technology and delivered as an oligo library pool with all 1,104 sequences in a single tube (10pmol lyophilized). The 10pmol pool of guide templates was resuspended in 100uL of nuclease-free water (final concentration: 1×10^−5^pM).

### Template Sequence

5’-TAATACGACTCACTATAG**-*20nt-seq*-**GTTTTAGAGCTAGAAATAGCAAGTTAAAATAAGGCTAGTCCGTTATCAACTTGAAAAAGTGGCACCGAGTCGGTGCTTTT-3’

The 20nt insert targeting sequences are described in Supplementary Table T2.

### Library amplification

The library was PCR amplified in a 20uL reaction using with 8μL of 5uM primer mix (T7_for [anchored]: 5’-CGCGCGTAATACGACTCACTATAG-3’; T7_rev:

5’-AAGCACCGACTCGGTGCC-3’), 1μL of the 1×10^−5^pM library stock, 1μL of DMSO (final 5%), and 10μL of Kapa 2X HiFi ReadyMix (Roche, KK2601) with the following parameters: 2min 95°C; 8x (20sec 95°C, 30sec 48°C, 30sec 72°C); 3min 72°C. Amplified libraries were then concentrated using a MinElute spin column (Qiagen, 28004).

### GuideRNA production

75ng of the amplified library was used as input for an RNA Synthesis Kit (NEB, E2050S) and concentrated using a Monarch RNA clean-up kit (NEB, T2040S). The IVT gRNAs were either kept on ice for use the same day or aliquoted and snap-frozen in liquid nitrogen.

### RNP complex assembly

50pmol of gRNA was combined with 25pmol of HiFi Cas9 Nuclease V3 (IDT, 1081060) in 1X CutSmart Buffer (NEB, B7204) at a final volume of 20μL. This was incubated at room temperature for 20 minutes, then stored on ice for use the same day.

### Cas9 Cleavage and Library Preparation

5μg of input DNA in 30μL of 1X CutSmart buffer (NEB, B7204), was dephosphorylated with 3μL of Quick CIP enzyme (NEB, M0508) for 10min at 37°C. The 20μL RNP complex mix, 1μL of 10mM dATP (Zymo, D1005), and 1μL of Taq DNA polymerase (NEB, M0267) were added for A-tailing of DNA ends. The sample was next incubated at 37°C for 20 minutes followed by 5 minutes at 72°C. Sequencing adaptors from the Oxford Nanopore Ligation Sequencing Kit (ONT, LSK109) were ligated to DNA ends using Quick Ligase (NEB, M2200) for 10 minutes at room temp. The sample was cleaned up using 0.3X Ampure XP beads (Beckman Coulter, A63881). Sequencing libraries were prepared by adding the following to the eluate: 25μL sequencing buffer (ONT, LSK109), 9.5μL loading beads (ONT, LSK109), and 0.5μL sequencing tether (ONT, LSK109). The sequencing reads were generated using Minion 9.4.1 flow cells.

### Cell lines

The GM12878 cell line was purchased from Coriell institute (CEPH/UTAH Pedigree 1463). The patient-derived PDAC cell lines were derived at Johns Hopkins, as described by Sjoblom et al^15^, with these lines previously described by Jones et al^3^. The Panc5.04 and Panc8.13 PDAC cell lines were originally developed by Dr. Elizabeth Jaffee^16^ and are available from ATCC; the A6L cell line was originally derived from a rapid autopsy study by Dr. Christine Iacobuzio-Donahue^17^. Cell lines were cultured in DMEM enriched with 10% FBS, 1% L-glutamine, and 1% penicillin-streptomycin; at 37°C in a humidified environment containing 5% CO_2_.

### Analysis of nanopore sequencing data

Basecalling was performed using guppy (Version 4) to generate FASTQ sequencing reads from electrical data. Reads were aligned to the human reference genome (hg19) using Minimap2 (v2.17). To calculate sequencing depth of coverage, BAM files were analyzed using Picard Tools (Broad Institute). Deletions were identified using the Sniffles SV caller (v2.0.2), with minimum length of 500bp and using the “--non-germline” option to identify rare variants^13^. SVs calls from Jones et al. were converted from hg17 alignments to hg19 alignments; Pa14C is Panc8.13, and Pa18C is Panc5.04, and Pa02C is A6L.

To quantify the reads generated per each gRNA, the number of forward strand reads started within 50nt downstream of the Cas9 cleavage site was counted for each gRNA. Statistical significance for comparing on-target and off-target scores of gRNAs was calculated using a two-tailed t-test (no assumption of equal variance). On-target gRNA scores were generated using either the Doench Score^7^, Zhang score^11^, or the online tool from Integrated Genome Technologies (IDT)^14^. The off-target score was from IDT^14^.

## Data availability

Sequencing data for this study can be retrieved from the Sequence Read Archive (SRA), under the BioProject ID PRJNA924081.

## Acknowledgments

This study was supported by the National Institutes of Health (grant R01HG009190).W.T. has two patents (8,748,091 and 8,394,584) licensed to ONT.

## REFERENCES

1. Nurk, S. et al. The complete sequence of a human genome. Science 376, 44–53 (2022).

2. Gilpatrick, T. et al. Targeted nanopore sequencing with Cas9-guided adapter ligation. Nat. Biotechnol. 38, 433–438 (2020).

3. Jones, S. et al. Core signaling pathways in human pancreatic cancers revealed by global genomic analyses. Science 321, 1801–1806 (2008).

4. Biankin, A. V. et al. Pancreatic cancer genomes reveal aberrations in axon guidance pathway genes. Nature 491, 399–405 (2012).

5. Norris, A. L., Workman, R. E., Fan, Y., Eshleman, J. R. & Timp, W. Nanopore sequencing detects structural variants in cancer. Cancer Biol. Ther. 17, 246–253 (2016).

6. Norris, A. L. et al. Transflip mutations produce deletions in pancreatic cancer. Genes Chromosomes Cancer 54, 472–481 (2015).

7. Doench, J. G. et al. Optimized sgRNA design to maximize activity and minimize off-target effects of CRISPR-Cas9. Nat. Biotechnol. 34, 184–191 (2016).

8. Farboud, B. & Meyer, B. J. Dramatic enhancement of genome editing by CRISPR/Cas9 through improved guide RNA design. Genetics 199, 959–971 (2015).

9. Chari, R., Mali, P., Moosburner, M. & Church, G. M. Unraveling CRISPR-Cas9 genome engineering parameters via a library-on-library approach. Nat. Methods 12, 823–826 (2015).

10. Konstantakos, V., Nentidis, A., Krithara, A. & Paliouras, G. CRISPR–Cas9 gRNA efficiency prediction: an overview of predictive tools and the role of deep learning. Nucleic Acids Res. 50, 3616–3637 (2022).

11. Hsu, P. D. et al. DNA targeting specificity of RNA-guided Cas9 nucleases. Nat. Biotechnol. 31, 827–832 (2013).

12. Sedlazeck, F. J. et al. Accurate detection of complex structural variations using single-molecule sequencing. Nat. Methods 15, 461–468 (2018).

13. Smolka, M. et al. Comprehensive Structural Variant Detection: From Mosaic to Population-Level. bioRxiv 2022.04.04.487055 (2022) doi:10.1101/2022.04.04.487055.

14. IDT Cas9 Tool. Integrated Genome Technologies OnlineCas9Tool https://www.idtdna.com/site/order/designtool/index/CRISPR_CUSTOM.

15. Sjöblom, T. et al. The consensus coding sequences of human breast and colorectal cancers. Science 314, 268–274 (2006).

16. Jaffee, E. M. et al. Development and characterization of a cytokine-secreting pancreatic adenocarcinoma vaccine from primary tumors for use in clinical trials. Cancer J. Sci. Am. 4, 194–203 (1998).

17. Mudali, S. V. et al. Patterns of EphA2 protein expression in primary and metastatic pancreatic carcinoma and correlation with genetic status. Clin. Exp. Metastasis 23, 357–365 (2006).

