## Supplementary Information for "IVT generation of guideRNAs for Cas9-enrichment Nanopore Sequencing"

Supplementary Tables:

Supplementary Table T1: All structural variants called in PDAC cell lines using Sniffles version 2 (Smolka, 2022). The highlighted rows indicate previously annotated SVs (Jones, 2008), with the notable exception of two highlighted chr9 deletions in Panc5.04 which were previously called as a single homozygous deletion.

Supplementary Table T2: The table shows the sequence for all gRNAs used in the study.

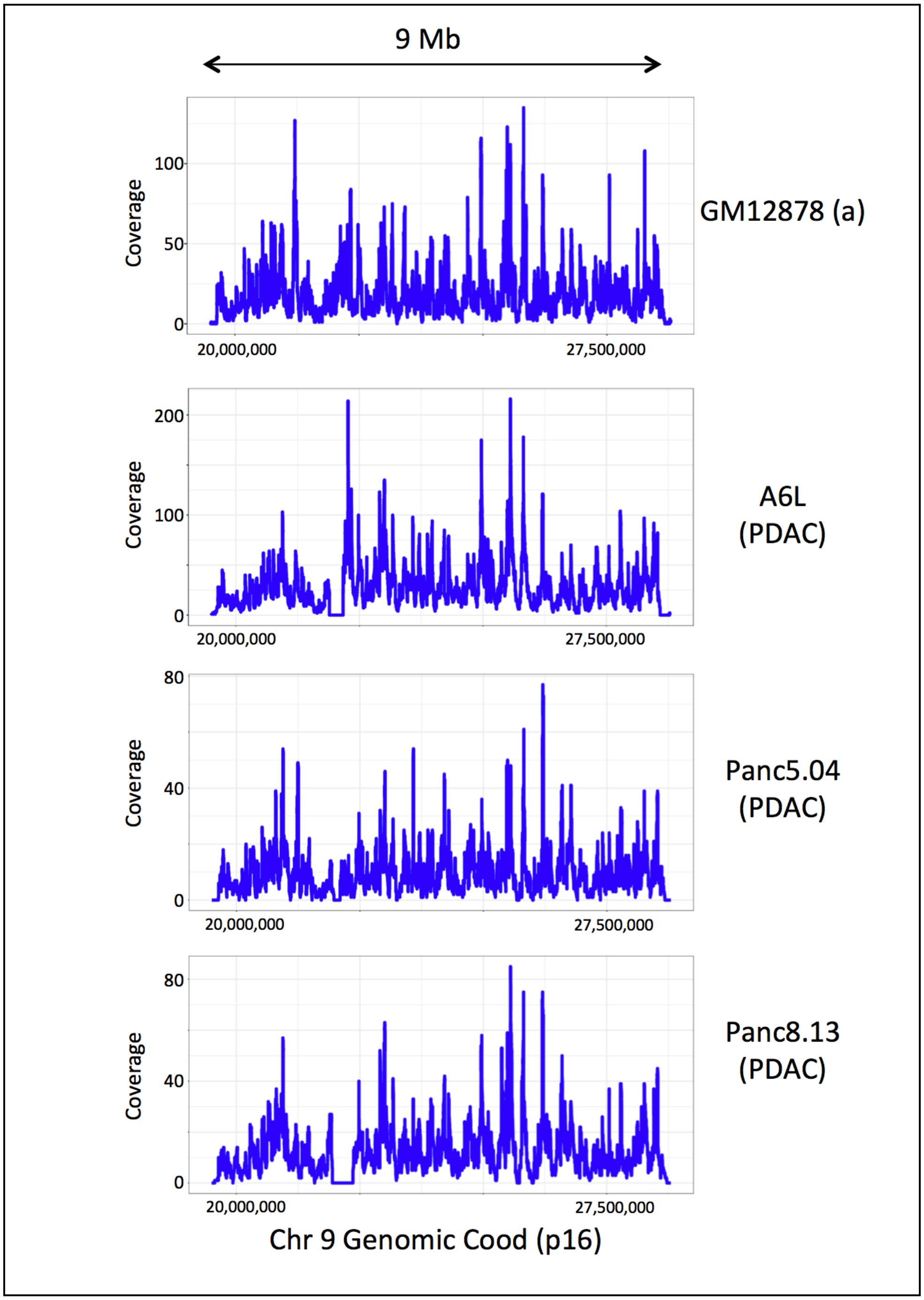


Supplementary Figure 1- Coverage in four cell lines across the entire enriched region of chr9.

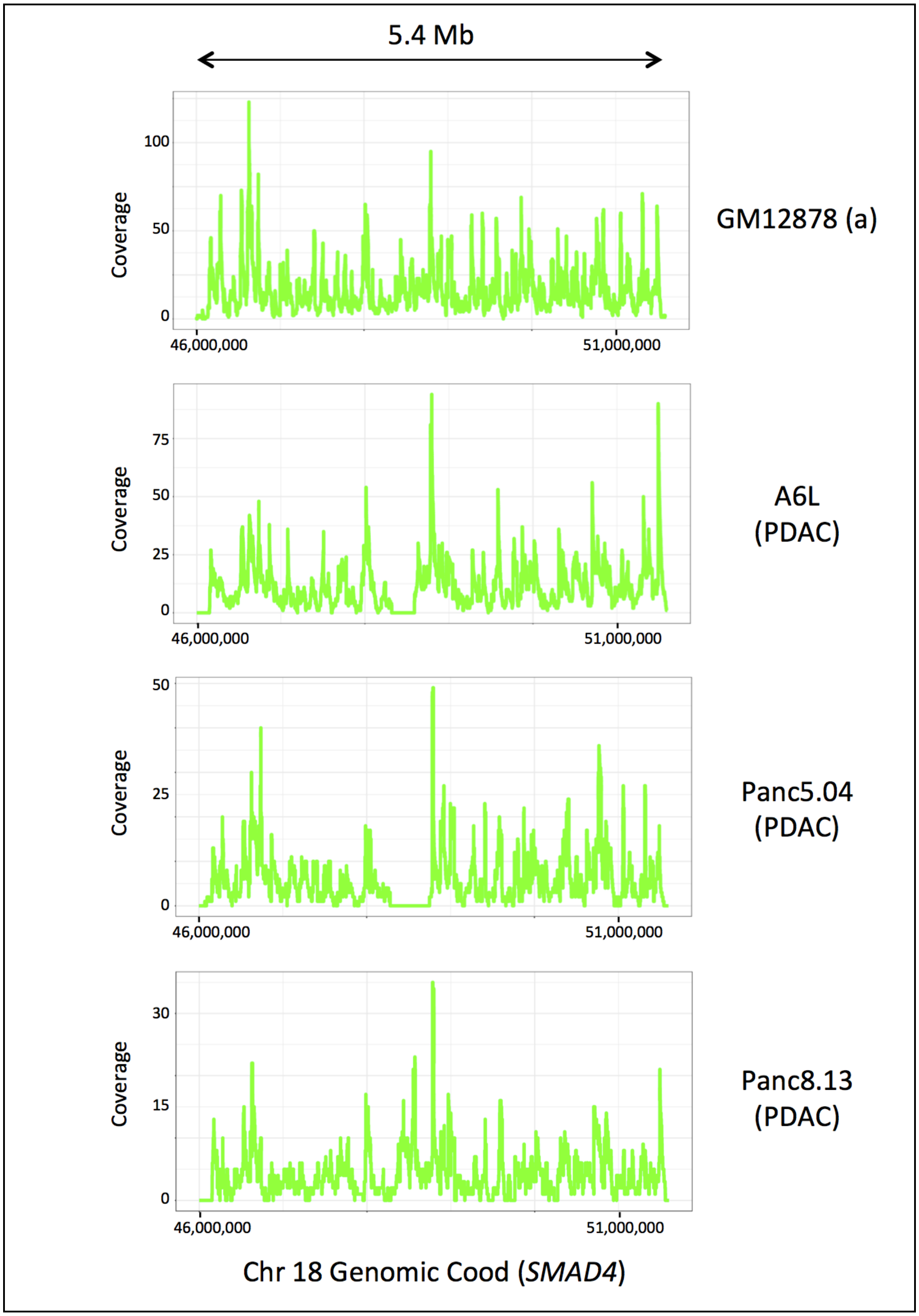


Supplementary Figure 2- Coverage in four cell lines across the entire enriched region of chr18.

| | SNP Array SV calls | | | | Targeted Nanopore SV calls | | | |
|---|---|---|---|---|---|---|---|---|
| | chr | L break | R break | deL size | chr | L break | R break | deL size |
| A6L | 18 | 48,321,014 | 48,588,407 | 267,393 | 18 | 48,319,051 | 48,589,278 | 270,227 |
| Panc5.04 | 18 | 48,326,378 | 48,746,375 | 419,997 | 18 | 48,279,930 | 48,749,687 | 469,757 |

Supplementary Figure 3 - Table comparing SV calls in two PDAC cell lines (Omni 2.5 SNP array data2 compared with Sniffles v2 calls on Cas9-enrichment nanopore data)
